# Tangled evolutionary history: genetically divergent taxa and hybrids characterise lantana invasions in Australia

**DOI:** 10.1101/2025.10.13.681924

**Authors:** Patricia Lu-Irving, Francisco Encinas-Viso, Eilish McMaster, Jason Callander, Michael Day, Johannes J. Le Roux

**Affiliations:** Research Centre for Ecosystem Resilience, Botanic Gardens of Sydney, Sydney, NSW, Australia; Centre of National Australian Biodiversity Research, CSIRO, Canberra, ACT, Australia; School of Life, Earth, and Environmental Sciences, University of Sydney, Sydney, Australia; Department of Primary Industries, GPO 267, Brisbane, QLD, Australia; School of Natural Sciences, Macquarie University, Sydney, NSW, Australia

**Keywords:** Australia, biological control, invasive species, *Lantana camara*, population genomics, species delimitation, weeds

## Abstract

**Aim:** to investigate population genomics and phylogeography in invasive lantana, including its taxonomy, spatial distribution, and patterns of morphological and genetic variation across the Australian continent.

**Location:** the main area invaded by lantana on the Australian continent, i.e., coastal and subcoastal eastern Australia from northern Queensland to southern New South Wales up to approximately 100 km inland, across 22 degrees latitude. The native range of lantana in the Americas was also represented.

**Methods:** we used DArTseq to generate genome-wide single nucleotide polymorphism (SNP) data for >600 individuals representing the native and Australian invaded ranges of lantana. We analysed data from >20,000 SNPs to identify distinct genetic clusters, and test the extent to which they corresponded with taxonomic descriptions and morphotype concepts used in lantana biological control. Genome sizes were estimated for a small representative subset of individuals using flow cytometry. We used MaxEnt to estimate habitat suitability for different lantana genetic clusters, and compared these predictions with observed patterns in biological control agent host-specificity.

**Results:** invasive lantana in Australia consisted of several divergent genetic clusters of most likely tetraploid individuals. Gene flow between genetic clusters was limited, consistent with the notion that multiple species introductions and/or hybridisation events were part of the invasion history. Two widespread, homogeneous genetic clusters were found to be dominant in Australia; two other genetic clusters with more limited distributions were identified with potential for future spread. Biological control agent host preferences were consistent with identified genetic clusters.

**Main conclusions:** treating invasive lantana as a single taxon may be counterproductive for effective management. Comprehensive taxonomic revision is needed to enable more precise identification of invasive taxa. Improved identification will support improved management, particularly if using biological control.

## INTRODUCTION

Invasive species are a major threat to biodiversity conservation (Kearney et al., 2019; Roy et al., 2024; Vilà et al., 2011). Accurate identification of invasive species is needed to effectively mitigate their impacts, e.g., for selection of biological control agents (Castillo et al., 2021; Pyšek et al., 2013), a highly cost-effective management approach (van Wilgen et al., 2012). Some factors that obscure the taxonomy of invasive species can also play a role in their ecological success, e.g., multiple introductions and admixture enabling rapid adaptation to new environments (Ellstrand & Schierenbeck, 2000; Le Roux & Wieczorek, 2009; Schierenbeck & Ellstrand, 2009). Therefore, understanding the taxonomy of invasive species is integral in understanding the processes contributing to invasion, and optimising management outcomes (Le Roux, 2021; Pyšek et al., 2013). A classic example of problematic taxonomic uncertainty in an invasive species is *Lantana camara* (Day et al., 2003a; Urban et al., 2011).

The *Lantana camara* species complex (*L. camara sensu lato*, hereafter referred to as “lantana”) has been listed among the world’s worst invasive species (Lowe et al., 2000). Invasive lantana populations are currently known from over 90 countries and have been documented to cause substantial economic, environmental, and human health impacts (e.g., AEC Group, 2007; Kearney et al., 2019; Silver & Carnegie, 2017; Syed & Guerin, 2004). In Australia, lantana is listed as one of 32 Weeds of National Significance (Thorp & Lynch, 2000) and has invaded millions of hectares over thousands of kilometres along the country’s eastern coast, spanning 22 degrees in latitude. Lantana has the greatest impact of any invasive plant on Australia’s threatened species (Kearney et al., 2019) and has been estimated to cause over AU$100 million in economic losses to agriculture annually (AEC Group, 2007).

Biological control of lantana has been implemented in many countries. However, despite 44 biological control agents released globally since 1902 (30 of these in Australia), lantana is still not considered to be under adequate control (Winston et al., 2014). Matching genotypes of invasive plants with their co-evolved specialist natural enemies can enhance the effectiveness of biological control programs (Goolsby et al., 2006), but this approach requires understanding the native origins of invasive populations (e.g., Jourdan et al., 2019). Thus, one of the main factors limiting successful biological control is the unresolved taxonomy of lantana (Day & Neser, 2000; Day et al., 2003b).

Invasive and cultivated forms of lantana are derived from a clade of up to 20 species naturally found from southern North America to southern South America (*Lantana* section *Lantana*; Lu-Irving et al., 2021; Sanders, 2006, 2012). While the name *Lantana camara* is routinely applied to invasive and cultivated members of section *Lantana*, their wide range of morphological variation does not align with Linnaeus’ (1753) concept of *L. camara*, nor with any other naturally occurring *Lantana* species (Sanders, 2006, 2012; Smith & Smith, 1982; Urban et al., 2011). This is consistent with a history of, and origin in, cultivation: different species of *Lantana* were extensively hybridised to create ornamental cultivars, particularly in the mid to late 19th century (Howard, 1969), when introductions to Australian gardens were likely to have taken place. Ornamental lantana was cultivated in multiple areas in Australia by 1850, and its invasiveness was noted by the end of the 19th century (Michael, 1987; Swarbrick, 1986).

Invasive lantana comprises extensive morphological diversity. Previous attempts to make sense of this variation include multiple, often competing, classification systems that range from formal names and descriptions (Munir, 1996; Sanders, 2006, 2012; Smith & Smith, 1982) to less formal working concepts (Day et al., 2003a, b; Heystek, 2006). Australian researchers used the informal concept of five “varietal groups” to identify putatively distinct host types for biological control (Day et al., 2003a). These groups (hereafter referred to as morphotypes) were based on aggregating the 29 varieties described by Smith & Smith (1982) according to flower colour: pink, red, pink-edged red, orange, and white (Fig. 1). Numerous biological control agents have failed to establish on some morphotypes, and establishment has been inconsistent with morphotype in Australia (Day et al., 2003a, b; Thomas et al., 2006). Thus, the five morphotype concept may inadequately reflect underlying variation in Australian lantana. The lack of a well-supported classification system means that most scientists, managers, and legislators take a broad taxonomic view of lantana, treating it as a single species. This view implies certain assumptions that are untested and may be invalid (e.g., all individuals are fully interfertile, morphological variants are not genetically divergent).

**Figure 1.**
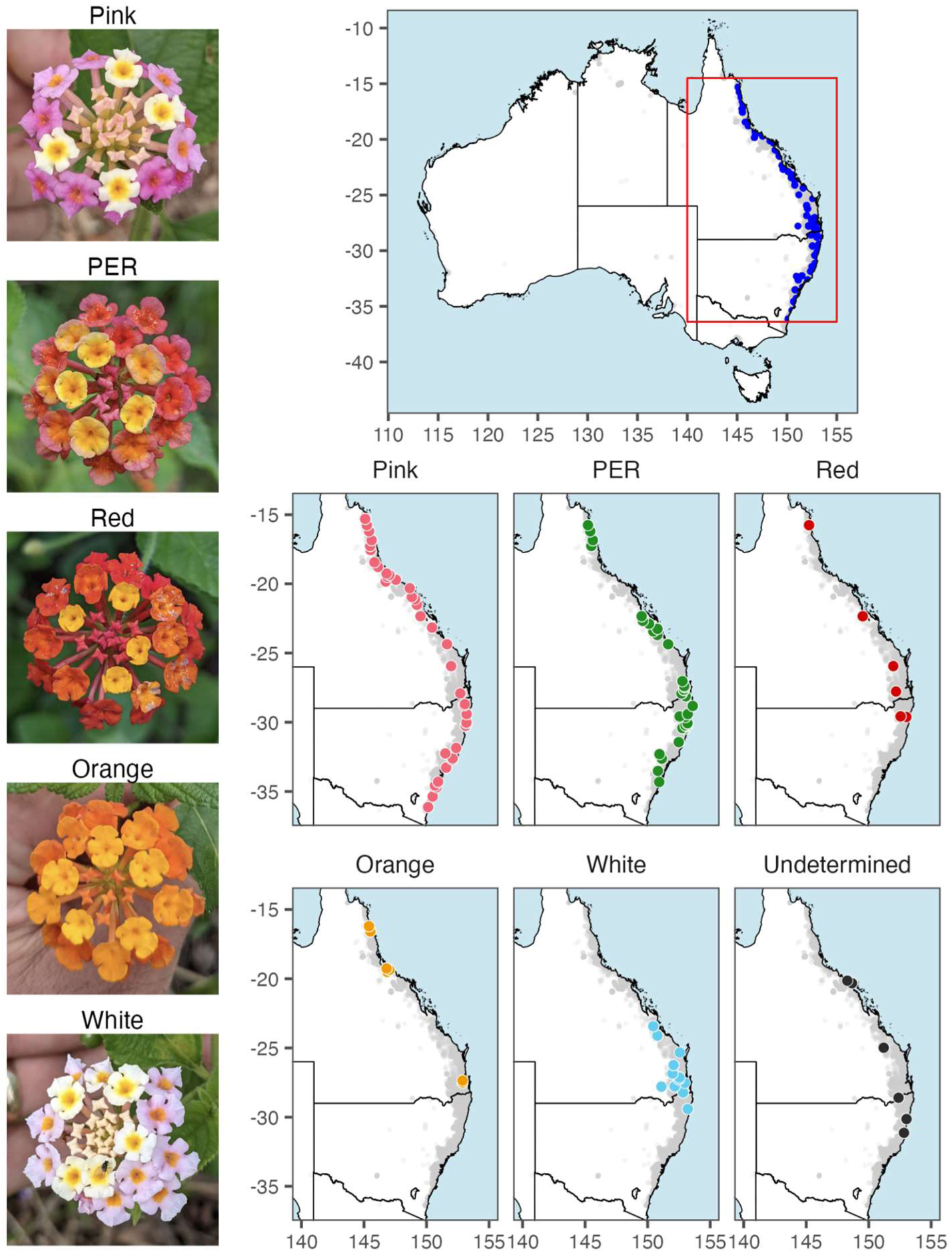
Flower morphotypes and distribution of lantana in Australia. Photographs of each of five flower colour morphotypes (varietal groups) recognised by the lantana biological control program in Australia, and maps showing their sampled distributions along the continent’s eastern coast. Photographs (top to bottom): inflorescences of individuals identified in the field as pink, pink-edged red (PER), red, orange, white morphotypes. All maps show pale grey points representing lantana occurrence records since 2000 from the Australasian Virtual Herbarium, documenting the current known distribution of lantana in Australia. Additionally plotted are points showing all locations sampled in this study (dark blue; top), and locations where samples were identified to morphotype or otherwise unable to be identified (six individually coloured maps; bottom).

A better understanding of the diversity of invasive lantana in Australia is urgently needed to better mitigate its impacts. In this study, we collected samples and data on an unprecedented scale to deliver new insights into the genetic composition and evolutionary history of lantana in Australia. We used DArTseq to discover thousands of single nucleotide polymorphisms (SNPs) across hundreds of lantana individuals from the native range in the Americas and the invaded range in Australia. Specifically, we aimed to characterise patterns of population genetic structure to test the hypothesis that *L. camara s.l.* in Australia represents a single biological species, versus the alternative hypothesis of multiple lineages separated by some degree of reproductive isolation, and with different ecological niches. Further, we hypothesised that genetic structure within lantana, if present, would align with the five flower colour morphotypes and with native-range provenance.

## METHODS

### Sampling

Wild plants belonging to *Lantana* section *Lantana* were sampled in the field: leaf tissue was collected and preserved in silica gel and/or by lyophilisation. In the invaded range along the eastern coast of Australia, a sample from each of six individuals (>10 metres apart where possible) was collected (with permission) at each site to maximise the capture of genetic diversity within and among sites. Where multiple morphotypes occurred at a site, six individuals per morphotype were sampled. Additional representation of both native and Australian ranges was achieved by sampling from specimens at Nagoya-compliant institutions Australian National Herbarium (CANB) and the Burke Museum Herbarium (WTU). In total, 537 individuals from 90 sites representing the invasion in eastern Australia and 166 individuals from the native range of *Lantana* section *Lantana* (Argentina, Belize, Bolivia, Brazil, Colombia, Costa Rica, Dominican Republic, Ecuador, Guatemala, Jamaica, Mexico, Peru, USA, Venezuela) were sequenced (Fig. 1, Table S1).

All five morphotypes described by Day et al. (2003a) were collected from multiple sites, with sampling proportions per morphotype of 36% pink, 34% PER, 11% white, 7% orange, 6% red, and 5% undetermined. We tried to sample as evenly as possible among the five morphotypes, but their relative abundance in the wild was highly uneven, i.e., pink was the most common flower colour observed, followed by PER, with other morphotypes relatively uncommon. This was broadly consistent with Day et al. (2003a), who recorded pink-flowered lantana at ∼75% of survey sites versus <20% of sites for other morphotypes.

### Sample identification

Smith & Smith (1982) described 29 varieties of *Lantana camara* in Australia, with multiple varieties of each flower colour. Distinctions between them were often subtle and/or subjective (e.g. relying on different shades or hues of colour) and sometimes based on overlapping ranges of measurements rather than categorical traits, making it challenging to identify plants to variety. Due to the difficulty using Smith & Smith’s (1982) treatment for identification, especially with pressed specimens (since the colour and shape of the corolla are both lost after pressing), we did not attempt to identify samples to varietal level. Instead, we made post-hoc comparisons between Smith & Smiths’ (1982) descriptions and field-collected specimens falling into different genetic clusters (see Results).

Sanders (2012) produced a comprehensive taxonomic revision of *Lantana* section *Lantana* in which he recognised 20 species (including 13 subspecies and five varieties), distinguished mainly by qualitative differences in leaf hair morphology. Identification of specimens this way was challenging due to the subtlety and subjectivity in interpreting the variation in this character, but some herbarium specimens were identified to species described by Sanders (2012) with the assistance of Sanders himself. Some specimens were morphologically determined to be non-hybrid individuals of the following nine species of *Lantana* section *Lantana* (the clade from which the invasive *L. camara s.l.* complex is derived): *L. bahamensis*, *L. camara*, *L. depressa*, *L. horrida*, *L. leonardiorum, L. nivea, L. scabrida, L. strigocamara, L. urticoides*. However, most specimens were determined to be putative hybrids (Table S1).

### Genotyping

Previous studies of *Lantana* section *Lantana* using DNA sequences from up to seven loci showed insufficient variation to resolve relationships among and within species (Lu-Irving & Olmstead, 2013; Scott et al., 1997; Watts, 2009). Studies using anonymous dominant genetic markers (e.g., RAPD, AFLP) had some success, but had insufficient data and/or taxon sampling to yield definitive insights (Scott et al., 1997; Watts, 2009). Genotyping-by-sequencing approaches, with their broader genomic coverage, were identified as a promising alternative (Lu-Irving et al., 2022). Here, we used the marker-discovery pipeline developed by Diversity Arrays Technology Pty Ltd (DArT), which has been successfully applied in various systems including non-model and polyploid plants (Kilian et al., 2012). Samples were sent to DArT in Canberra for DNA extraction and DArTseq analysis following established procedures (Jaccoud et al., 2001; Rossetto et al., 2018). DNA was extracted and genome-wide marker data were generated via DArTseq, provided as biallelic SNP genotype matrices.

### Population genetic diversity and structure of Australian lantana

To better understand patterns of genetic diversity and relationships among invasive lantana populations in eastern Australia, a dataset consisting of only samples collected in the states of Queensland (Qld) and New South Wales (NSW) was analysed. All analyses were conducted in the R statistical environment (R v4.1.0; R Core Team, 2021). To ensure data quality, loci with reproducibility <98% or missing data >80% were removed. Fixed loci were excluded, and only one SNP per DArT tag was retained. Samples with >80% missing data were discarded. Prior to conducting Principal Component Analysis and calculating diversity statistics, loci were further filtered to retain only those with a minor allele frequency (MAF) of ≥2%.

Principal Component Analysis (PCA) was conducted using the *adegenet* package (v2.1.10; Jombart, 2008). To identify genetic clusters, t-distributed stochastic neighbour embedding (t-SNE) was performed. The genotype data was converted to a Euclidean distance matrix, and the t-SNE algorithm run using the *Rtsne* package (v0.17) with parameters set to a perplexity of 15, and a theta of 0. Clusters were subsequently identified by applying hierarchical density-based spatial clustering (HDBSCAN) using the *dbscan* v1.1-12 package (Hahsler et al., 2019; Scitovski & Sabo, 2020) with a minimum points parameter of 5. Clusters containing fewer than 10 samples were removed, as were clusters that were not monophyletic (as assessed using Unweighted Pair Group Method with Arithmetic mean (UPGMA) clustering on Euclidean distances).

To evaluate patterns of genetic structure, ancestry estimation was conducted using sparse non-negative matrix factorization (sNMF) implemented in the *LEA* package (v3.14.0; Frichot & François, 2015) across a range of K values (1–20) with default settings.

Pairwise fixation indices (*F*_ST_) were calculated using the relative beta estimator (Weir & Hill, 2002) with the *SNPrelate* package (v1.20.1; Zheng et al., 2012). Genetic clusters were identified using HDBSCAN (see Results). A linear model was used to test if *F*_ST_ differences between sites were influenced by cluster membership (intra-cluster, inter-cluster, or unclustered sites) and pairwise geographic distance. The analysis was performed using the *stats* package (v4.3.1).

Genetic diversity was assessed using basic diversity statistics: observed heterozygosity (*H_O_*) and inbreeding coefficient (*F_IS_*) (Keenan et al., 2013).

#### Identification of putative hybrids

Plants at two NSW sites (Korora and Nana Glen) were found to be morphologically intermediate between pink and PER, sympatric with putatively non-hybrid plants. We compared observed heterozygosity and fixed loci between putative hybrids and non-hybrids at these two sites and at two nearby sites where only non-hybrid plants occurred.

### Relationships among Australian and native-range lantana

To infer relationships among samples from eastern Australia and the native range, we used two distance-based approaches. Loci were filtered as described above.

A phylogenetic network was constructed in SplitsTree v4.18.2 (Huson & Bryant, 2006) to assess clustering and hybridisation patterns, using a Euclidean distance matrix and the *RSplitsTree* v0.1.0 package. The network was plotted in R using *tanggle* v1.8.0 (Schliep et al., 2024). Phylogenetic relationships were also assessed using UPGMA, which bypasses assumptions of model-based approaches, which are problematic in this case due to sparse SNPs, lack of reference genomes, and the potential effects of hybridisation. A Hamming distance matrix was used to generate UPGMA dendrograms, with bootstrap support (10,000 replicates) calculated using the *phangorn* v2.11.1 package (Schliep, 2011) to assess the topology’s robustness.

### Species distribution modelling

To understand how different environmental variables influence the distribution of invasive lantana in Australia, we analysed sample occurrence data to predict habitat suitability in Australia of different genetic clusters (see Results).

Habitat suitability was estimated with species distribution models using the maximum-entropy (MaxEnt) method based on the environmental variables of sample collection coordinates. Species distribution models were constructed using MaxEnt v3.4.1 software (Phillips et al., 2017) and the *dismo* v1.3–3 (Hijmans et al., 2024) R package using all specimen records as training data, 21 environmental variables as predictor layers at 1 km^2^ resolution, and 10,000 background points.

Specimen records corresponded with field collection coordinates for samples falling into identified genetic clusters (see Results; Fig. S2).

The environmental predictors used for the MaxEnt models were 15 bioclimatic variables and six soil and landform variables. Bioclimatic variables were: temperature seasonality (CV; Bio04), maximum temperature of warmest month (°C; Bio05), minimum temperature of coldest month (°C; Bio06), mean temperature of warmest quarter (°C; Bio10), mean temperature of coldest quarter (°C; Bio11), annual precipitation (mm; Bio12), precipitation seasonality (CV; Bio15), precipitation of driest quarter (mm; Bio17), annual mean moisture index (Bio28), moisture index seasonality (CV; Bio31), mean moisture index of wettest quarter (Bio32), annual total actual evapotranspiration (mm; EAA), minimum monthly potential evaporation (mm; EPI), minimum monthly atmospheric water deficit (mm; WDI) and maximum monthly atmospheric water deficit (mm; WDX). Soil and landform variables were: soil clay content 0–30 cm (%; CLY), soil sand content 0–30 cm (%; SND), total soil nitrogen 0–30 cm (%; NTO), total soil phosphorus 0–30 cm (%; PTO), soil bulk density 0–30 cm (g/cm3; BDW) and topographic wetness index (TWI3S). Data (9 s resolution) are available from the CSIRO Data Access Portal at https://doi.org/10.25919/5dce30cad79a8 (Bio4-Bio32), https://doi.org/10.4225/08/5afa9f7d1a552 (EAA, EPI, WDI and WDX) and https://doi.org/10.4225/08/5b285fd14991f (CLY, SND, NTO, PTO, BDW and TWI3S). The Cloglog transform as a continuous index of habitat suitability was selected as the output format (Phillips et al., 2017). The maximum of Cohen’s Kappa (K, which determines the optimal threshold for statistical discrimination of presence-absence; Cohen, 1960), the maximum training sensitivity plus specificity (MTSS, equivalent to finding the point in the receiver operator characteristic (ROC) curve with a tangent slope of 1; Cantor et al., 1999) and the area under the ROC curve (AUC) were applied to test model performance.

### Genome size estimation

Polyploidy is common in native and invasive populations of *Lantana* Sect. *Lantana* (Sanders, 1987; Spies, 1984) and variation in ploidy can provide evidence of hybridisation. Flow cytometry was performed to measure and compare genome size as an indicator for ploidy variation among 14 individuals of the pink and PER morphotypes from five sites. *Glycine max* was used as an internal standard. Whole nuclei were extracted from fresh leaf tissue following standard protocols. Briefly, up to 1 cm^2^ of leaf tissue was chopped in Sysmex Cystain PI Abs. P extraction buffer, and filtered before staining with Sysmex Cystain PI Abs. P staining buffer and incubating at room temperature for 30-60 min. Samples were analysed using a BD Accuri flow cytometer.

## RESULTS

### Genotyping

Of the 703 samples of lantana submitted for SNP discovery, genotype data were returned for 687 samples; 16 samples did not yield usable data (Table S1).

Following processing and filtering of SNP matrices, two datasets were generated for further analysis: (1) 512 eastern Australian samples by 10,078 loci for population genetic analyses (4,986 loci after filtering with MAF ≥ 2%); (2) 524 eastern Australian samples and 128 native range samples (652 total samples) by 25,751 loci for phylogenetic inference (6,462 loci after filtering with MAF ≥ 2%).

### Population genetic diversity and structure of Australian lantana

Principal Component Analysis (PCA) revealed patterns among Australian lantana samples that partially aligned with morphotypes (Fig. 2a), but did not clearly separate them. Dimensionality reduction using t-SNE produced more distinct subgrouping (Fig. 2b). Clustering on t-SNE with HDBSCAN identified seven genetic clusters (groups with ≥10 individuals, monophyletic in UPGMA), labeled A–G (Fig. 2c–d). Of 512 Australian samples, 62% were assigned to one of these clusters (Table 1). Flower colour morphotypes were consistent within clusters but not diagnostic, because two morphotypes each spanned two clusters. The pink morphotype (36% of samples) encompassed clusters A and F, while the orange morphotype included clusters E and G. The t-SNE with HDBSCAN method groups samples by genetic similarity, requiring a minimum density of similar individuals.

**Figure 2.**
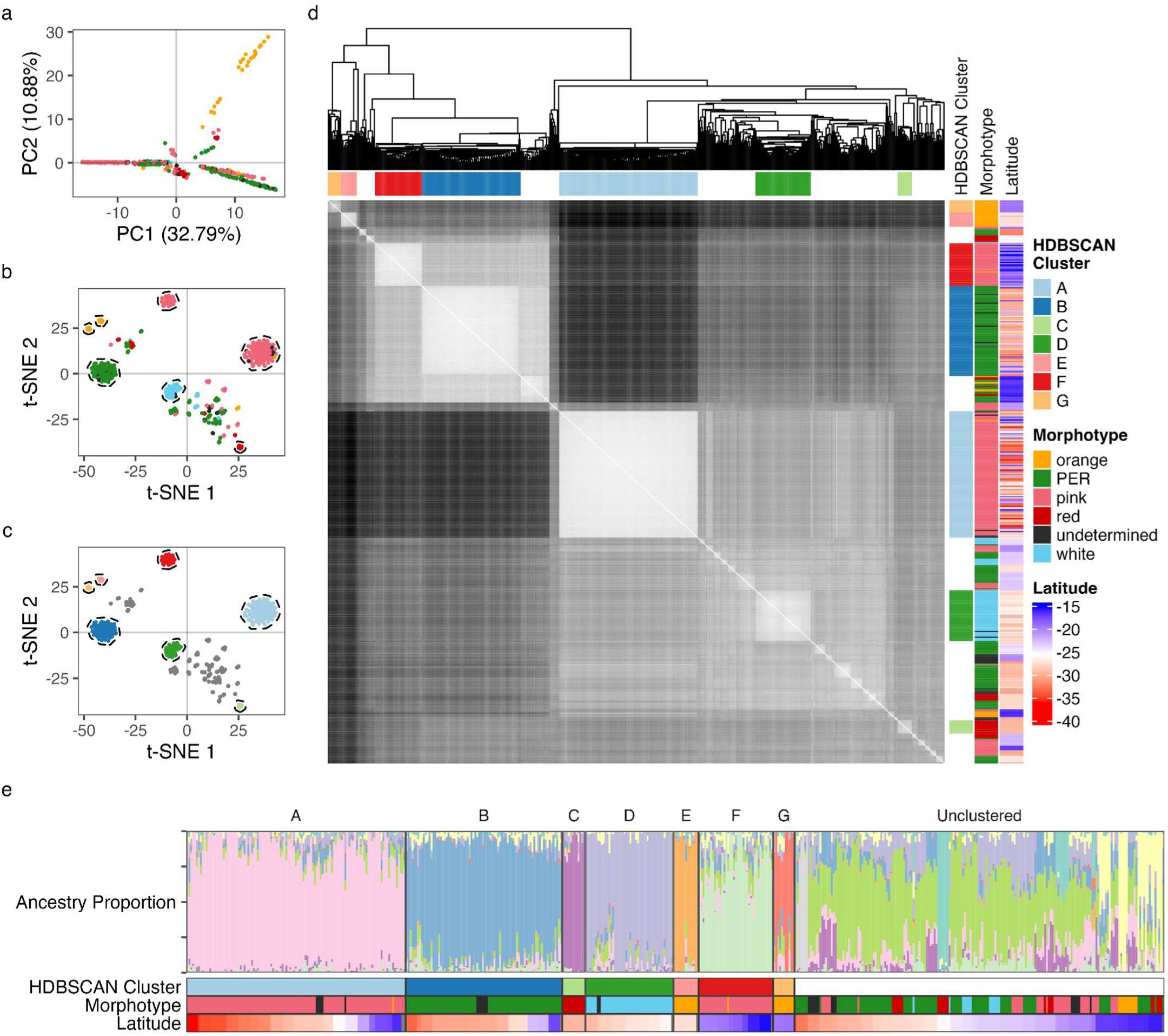
Cluster analyses of Australian lantana. Analyses of genetic clustering among 512 wild *Lantana camara s.l.* samples from eastern Australia: (A) plot of first two Principal Components identified by PCA with points coloured by morphotype; (B) t-SNE plot of samples, coloured by morphotype, with clusters identified by HDBSCAN circled. (C) t-SNE plot of samples, coloured by HDBSCAN clusters. (D) Heatmap of Euclidean distances between samples, ordered by UPGMA clustering (dendrograms shown). Adjacent annotations indicate HDBSCAN clusters, morphotypes, and collection latitudes. (E) LEA snmf barplot of samples (vertical bars) showing proportion/ancestry coefficient for each of K = 11 inferred ancestral populations (coloured vertical bars), with samples grouped by HDBSCAN clusters, with morphotype and latitude annotations below.

**Table 1.** Morphotype distribution across genetic clusters. Morphotype distribution across genetic clusters in invasive *Lantana camara* in eastern Australia. The table shows the number of samples for each morphotype (Pink, PER, Red, Orange, White, Undetermined) within seven genetic clusters (A–G) and an unclustered group. Each cluster is associated with a dominant morphotype, a descriptive variety name from Smith & Smith 1982, and the origin of the closest native-range relative where known. Percentages reflect the proportion of samples in each cluster relative to the total dataset (n = 512).

|  |  | Morphotype |  |  |  |  |  | Cluster n | Variety | Origin of closest native-range relative |
| --- | --- | --- | --- | --- | --- | --- | --- | --- | --- | --- |
|  |  | Pink | PER | Red | Orange | White | Undetermined |  |  |  |
| Cluster | A | 109 | 0 | 0 | 1 | 0 | 5 | 115 [22%] | "Common Pink" | Brazil |
|  | B | 0 | 76 | 0 | 0 | 0 | 6 | 82 [16%] | "Common PER" | Mexico |
|  | C | 0 | 0 | 12 | 0 | 0 | 0 | 12 [2%] | "Oblong Red" | - |
|  | D | 0 | 0 | 0 | 0 | 44 | 2 | 46 [9%] | "Helidon White" | - |
|  | E | 0 | 0 | 0 | 13 | 0 | 0 | 13 [3%] | "True Orange" | - |
|  | F | 38 | 0 | 0 | 1 | 0 | 0 | 39 [8%] | "Townsville Red-centred Pink" | Mexico |
|  | G | 0 | 0 | 0 | 11 | 0 | 0 | 11 [2%] | "Townsville Prickly Orange" | - |
|  | Unclustered | 39 | 96 | 21 | 10 | 13 | 15 | 194 [38%] |  |  |
| Morphotype n |  | 186<br>[36%] | 172<br>[34%] | 33<br>[6%] | 36 [7%] | 57<br>[11%] | 28<br>[5%] | 512 |  |  |

Samples that were genetically intermediate, highly variable, or located at cluster edges lacked sufficient similarity to cluster. Unclustered samples (38% of samples) included all morphotypes and were distributed throughout the geographic area sampled.

The seven genetic clusters defined by t-SNE and HDBSCAN ranged from common and widespread to infrequent and geographically restricted. They were strongly genetically isolated regardless of geographic proximity (Fig. 3) and varied in within-cluster genetic diversity. Each cluster corresponded to a variety described by Smith & Smith (1982). Cluster information is summarised in Table 1; see also Fig. 2d (within-vs. between-cluster genetic similarity), Fig. S2 (cluster distributions), Fig. S3 and Tables S3–4 (heterozygosity). Cluster A (n=115, pink) was widespread from northern Qld to southern NSW, matching the description and distribution of “Common Pink”. Cluster A had low heterozygosity (Ho 0.015 ± 0.004) and high genetic similarity among samples. Cluster B (n=82, PER), mainly in NSW but also occurring in Qld, matched “Common Pink-edged Red,” with similarly low heterozygosity (0.003 ± 0.003) and high similarity among samples. Cluster C (n=12, red) comprised only two sites in northern NSW, matched “Oblong Red” and had high heterozygosity (0.153 ± 0.02). Cluster D (n=46, white) matched “Helidon White,” and also had high heterozygosity (median *H*o: 0.12 ± 0.022). Smith & Smith (1982) recorded this variety only in southeast Qld; we found it abundant in northern NSW as well. Cluster E (n=13, orange), limited to southeast Qld, matched “True Orange”, with high heterozygosity (0.151 ± 0.034). Cluster F (n=39, pink), limited to northern Qld, matched “Townsville Red-centred Pink”, with low heterozygosity (0.019 ± 0.008). Cluster G (n=11, orange), also in northern Qld, matched “Townsville Prickly Orange” with variable heterozygosity (0.06 ± 0.045).

**Figure 3.**
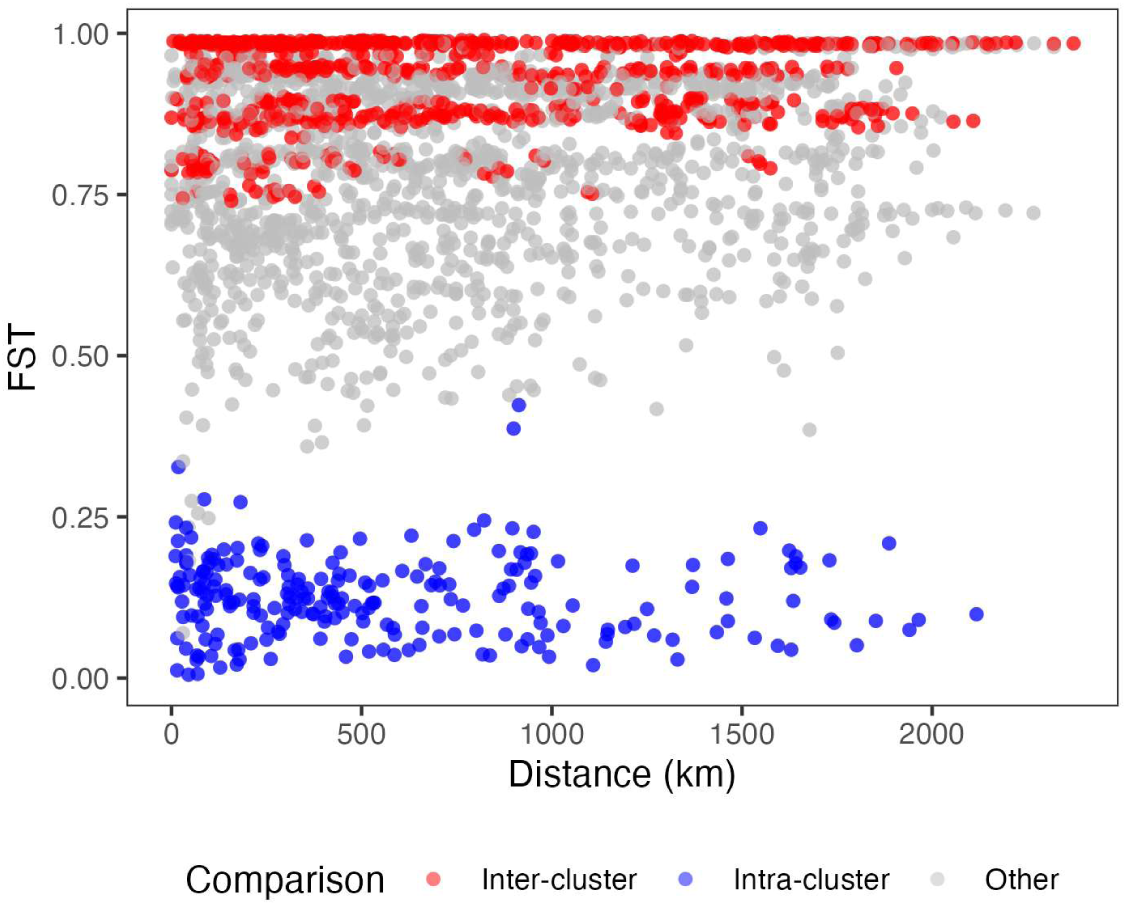
Pairwise F_ST_ and geographic distance between sites. Pairwise *F*_ST_ values between lantana collection sites in eastern Australia. Pairwise comparisons within the same HDBSCAN cluster are coloured blue (intra-cluster), comparisons between different HDBSCAN clusters are red (inter-cluster), and comparisons involving ungrouped sites are grey (other). All sites consist exclusively of samples from a single HDBSCAN cluster or ungrouped samples. Linear model results showed that both genetic cluster membership (inter-cluster, intra-cluster, other) and geographic distance had statistically significant effects on pairwise *F*_ST_ (p < 2e-16 for all variables).

Results of LEA snmf analysis were used to estimate ancestry and further explore genetic structure. These results were not used to define genetic clusters, but were compared with the results of cluster analysis. Ancestry patterns largely aligned with HDBSCAN clusters, and K-values >7 revealed substructure within the “unclustered” group (Fig. 2e, Fig. S4); since these additional putative genetic clusters had <10 samples they were not captured by HDBSCAN. Across tested K-values, K>10 best fit the data (minimised cross-entropy; Fig. S4). Figure 2e shows K=11 to best visualise patterns in genetic structure, but this should not be interpreted as the “true” number of ancestral lineages given the exploratory nature of the analysis (Meirmans, 2015) and the possibility of unsampled lineages.

Pairwise *F*_ST_ values between eastern Australian sites (Fig. 3) revealed strong differentiation between genetic clusters. These results were not used to define clusters but to assess gene flow within and between them. Estimated *F*_ST_ within clusters was consistently low (<0.25), indicating little differentiation regardless of geographic separation. Between clusters, *F*_ST_ was consistently high (>0.75), indicating limited gene flow even when sites were close or sympatric. Estimates involving unclustered sites showed variable *F*_ST_ (but typically >0.25), suggesting at least some isolation between sites and from defined genetic clusters. While these estimates may appear high compared with other datasets (*F*_ST_ ∼0.25 has generally been interpreted as strong differentiation, vs relatively weak in this study), this is typical for analyses of DArT data (McMaster et al., 2024). The linear model showed that both cluster membership and geographic distance significantly influenced pairwise *F*_ST_ (p < 2e-16 for all variables). Pairwise *F*_ST_ was lowest within clusters (mean 0.112), highest between clusters (mean 0.904), and high for unclustered sites (mean 0.767). Distance was positively correlated with *F*_ST_, but with a very small effect size (+0.00003 per km). The model explained 78.18% of the variation in pairwise *F*_ST_ (p < 2.2e-16, F-statistic: 5932 on 3 and 4966 DF).

#### Identification of putative hybrids

Morphologically-determined putative hybrids showed higher individual heterozygosity compared to co-occurring individuals identified as pink or PER, as would be expected for hybrids compared with their parents. On PC1 of the PCA, morphologically intermediate individuals clustered between the two genetic clusters corresponding with the pink and PER individuals. Additionally, putative hybrids were predominantly heterozygous at loci that were reciprocally fixed in pink and PER individuals (Fig. S5). These findings supported the hypothesis that morphologically intermediate individuals were hybrids resulting from interbreeding between different genetic clusters.

### Relationships among Australian and native-range lantana

The seven genetic clusters identified from analysis of the Australian dataset were recovered as well-defined groups in the phylogenetic network and as clades in the UPGMA tree (Fig. 4). The phylogenetic network displayed extensive webbing between groups, indicating reticulate evolution (e.g., historic hybridisation). Similarly, the UPGMA topology had poor bootstrap support at deeper nodes, reflecting lack of resolution (to be expected when bifurcating trees are inferred to represent non-bifurcating, i.e. reticulate, evolutionary relationships).

**Figure 4.**
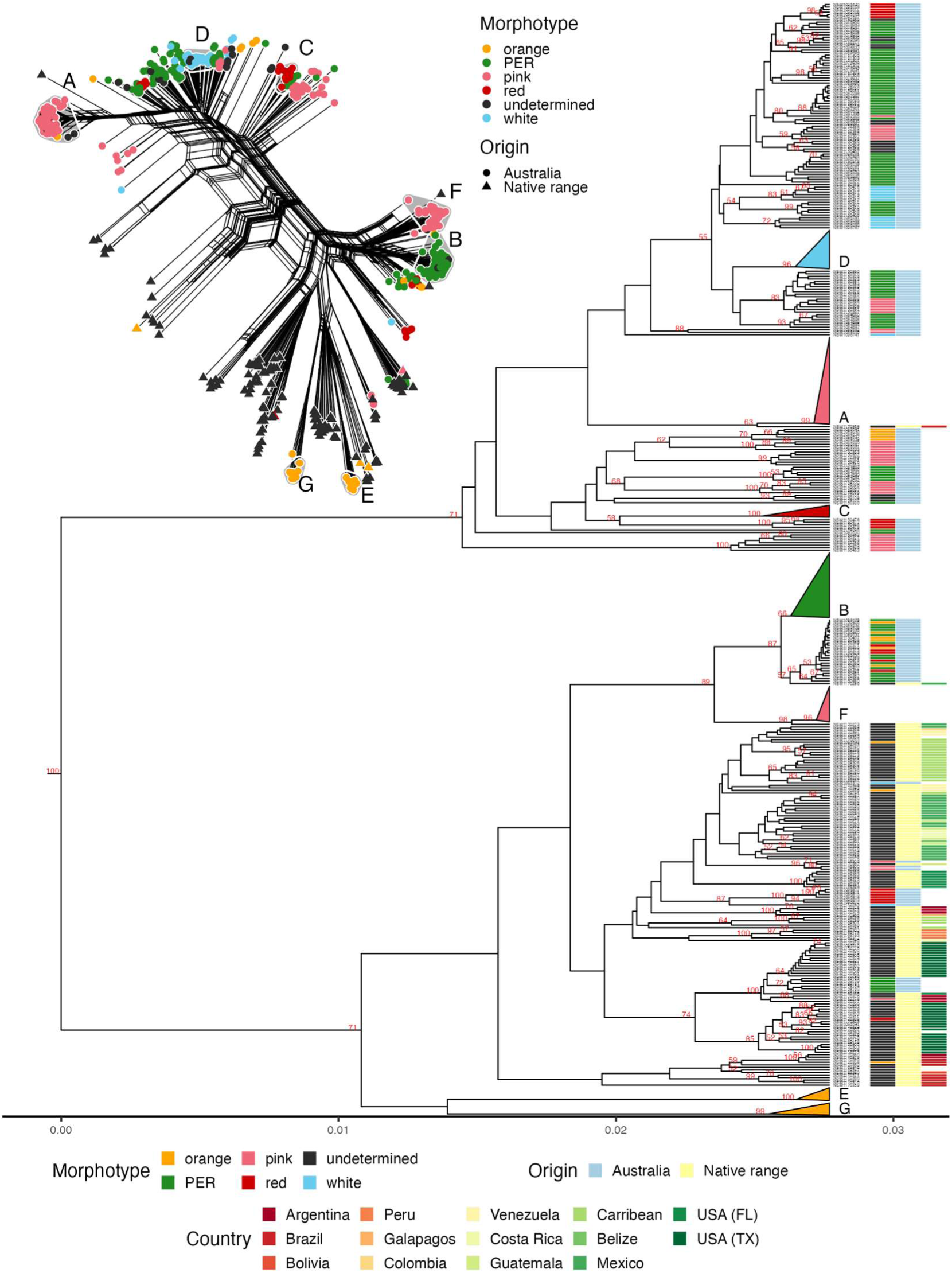
Phylogenetic relationships of Australian and native range lantana. Phylogenetic network (top left) and UPGMA tree depicting the Hamming distance between samples of *L. camara s.l.* from eastern Australia and native-range members of *Lantana* section *Lantana*. In the network, taxa are coloured according to morphotype and their origins are indicated by shape (circle = Australia, triangle = native range). Australian samples corresponding with HDBSCAN clusters are shaded with adjacent cluster labels A-G. The UPGMA tree shows bootstrap values >50; clades corresponding with Australian HDBSCAN clusters are collapsed and labelled A-G. Morphotype and origin are indicated for tips representing unclustered individuals.

Most native range samples did not group closely with any Australian samples, including those identified as putatively pure species, and Australian genetic clusters were not closely related to any of the native-range taxa analysed. However, cluster A was closely related to a single unidentified sample from southern coastal Brazil, cluster B was related to a single sample from Mexico, and cluster F was related to a different sample from Mexico. In contrast, clusters D and C showed no close relationship to any native range samples. Clusters E and G nested closely with native-range samples in the phylogenetic network, but no sister relationships with specific taxa or samples were recovered in the UPGMA tree.

Several unclustered Australian samples showed closer relatedness to native range samples than to the majority of Australian samples, e.g., six PER samples from Mount Thorley, NSW, were closely related to samples from Texas, USA; three pink samples from near Townsville, Qld were closely related to a Guatemalan sample; one white sample from near Emerald, Qld, was closely related to samples from the Dominican Republic. Additionally, six red samples from Booubyjan, Qld, and one white sample from Brisbane, Qld, clustered more closely with native-range samples than other Australian samples.

### Species distribution model

Model performance evaluated using the area under the receiver operating characteristic curve (AUC) was high for all genetic clusters (AUC ≥ 0.99; Table S2). Two widespread clusters were found to already occupy most of their predicted suitable habitat (clusters A and B); three clusters with more limited distributions also occupied most of their suitable habitat (clusters C, E, G), and two had much larger predicted suitable habitat than their sampled distributions (clusters D and F).

The most important variables determining the suitability of habitats for the seven genetic clusters were Annual mean moisture index (B28), Moisture index seasonality (B31) and mean moisture index of wettest quarter (B32) (Table S2).

### Genome size estimates

Genome size estimates obtained via flow cytometry ranged from 6.08 to 6.89 pg (2C), with 2C DNA content of the pink morphotype being on average 0.27 pg greater than that of the PER morphotype (Fig. S1). These estimates are broadly consistent with previously published genome size estimates in tetraploid *Lantana* Sect. *Lantana* (i.e., ∼6.3 Brooks Parrish et al., 2021; Deng et al., 2017).

## DISCUSSION

### Australian lantana consists of multiple, divergent genetic clusters

The invasive lantana complex in Australia is genetically diverse, comprising at least seven distinct genetic clusters. Limited gene flow between them suggests these clusters are largely reproductively isolated, contradicting the view of Australian lantana as a single morphologically variable species (e.g., Johnson, 2007; Munir, 1996). Flower colour morphotype was consistent within clusters, but failed to diagnose them, e.g., most pink-flowered samples (59%) belonged to cluster A, but 20% fell into cluster F, and 21% were unclustered.

Clusters A and B, the two most widespread and abundant in Australia, extended 2,250 km and 1,870 km along the eastern coast, spanning ∼19 and ∼16 degrees of latitude respectively. They exhibited the two most common morphotypes (pink and PER). Because our sampling was biased towards representing rarer morphotypes, the proportion of A and B in our dataset (∼40%) underestimates their actual abundance; we suggest that they comprise >50% of all invasive lantana in Australia (consistent with Day et al., 2003a). Clusters A and B were highly differentiated (*F*_ST_ ∼1) even in sympatry, indicating reproductive isolation, though occasional hybrids were observed (Fig. S5). Genome size estimates in both clusters were comparable with tetraploid *Lantana* Sect. *Lantana* (6.08–6.42; Brooks Parrish et al., 2021; Deng et al., 2017; Fig. S1), but a slight difference in average genome size may reflect differing DNA content and arrangement.

Genetic diversity within clusters A and B was remarkably low. No differentiation was detected among sites within the same cluster, even across large distances; e.g., *F*_ST_ between cluster A populations from Daintree Tea (northern Qld) and Norah Head (central NSW) was ∼0 despite >2,000 km separation between them. Clusters A, B, and F also showed distinctly low heterozygosity. Low heterozygosity and lack of isolation by distance are consistent with recent range expansion following a genetic bottleneck involving inbreeding (e.g., self-fertilisation), though they may also at least partially reflect SNP-calling biases in polyploid, multispecies datasets. In either case, this study lays a foundation for further demographic analyses focused on individual genetic clusters.

Almost 40% of Australian lantana samples were unclustered (showing no clear affiliation with the seven genetic clusters), occurring across the full invaded range in Australia. Some had mixed ancestry, while others likely represented additional clusters undetected due to insufficient sampling (<10 samples). For example, unclustered samples included hybrids between clusters A and B (Fig. S5), as well as samples comprising ancestral populations at K=11 (Fig. 2e). Thus, Australian lantana can be characterised as a mosaic of distinct genetic clusters, from common and widespread to rare and highly localised, interspersed with inter-cluster hybrids. This complexity explains why the taxonomy of lantana has been (and will continue to be) challenging.

Effective regulation and management of invasive species depends on accurate taxonomy that meaningfully describes them and separates them from other taxa (Pyšek et al., 2013). Our results did not support the concept of Australian lantana as a single species (i.e., individuals freely interbreeding resulting in genomic isolation by distance), nor did they support Sanders’ (2006, 2012) concept of a hybrid swarm descended from three principal parental species. Instead, our findings aligned best with the taxonomic treatment of Smith & Smith (1982), who described 29 distinct taxa. While we did not sample all 29, or identify all our samples using this treatment, the seven genetic clusters we identified were clearly equivalent to seven of the taxa described. These taxa were described as varieties, implying limited or weak evolutionary divergence, but they might be more suitably recognised as subspecies or even species, given the extent of genetic differentiation. Pending such a revision, we hereafter refer to the seven Australian genetic clusters identified in this study by their corresponding varietal names *sensu* Smith & Smith (1982).

### Cryptic patterns of provenance and evolutionary history

Our dataset included non-hybrid specimens of nine native-range *Lantana* sect. *Lantana* species, but none of the seven Australian clusters were closely related to them. Most native-range samples, including all non-hybrid specimens, fell into two clades containing only a few unclustered Australian samples. Three Australian genetic clusters were related to single, putatively hybrid, native-range samples from Brazil (cluster A) and Mexico (clusters B, F). No native-range sister lineage was identified for the remaining four clusters (C, D, E, G). The identification of native-range provenances of three Australian lantana varieties based on the results presented here should be treated with caution. However, we hypothesise based on these preliminary results that “Common Pink” (cluster A) was derived from plants native to Brazil, while the progenitors of “Common Pink-edged Red” (cluster B) and “Townsville Red-centred Pink” (cluster F) were native to Mexico. Testing this hypothesis will require further investigation with potentially important implications for improving lantana biological control (discussed below).

Our inability to identify the native-range provenance of Australian lantana may be due to limited sampling of the Americas. *Lantana* sect. *Lantana* comprises 20 species (some rare or even extinct in the wild; Sanders, 2006, 2012) from the southern USA to northern Argentina, but only nine were included in our analyses. Also, the garden origin of invasive lantana is potentially confounding: the introduction and naturalisation of multiple hybrid cultivars, possibly with further introgression taking place in the wild, would make inferring the ancestral range of present-day invasive populations difficult, if not impossible. We focused on characterising genetic diversity in Australian invasive lantana, rather than on comprehensive phylogenetic reconstruction of *Lantana* sect. *Lantana*, limiting the broader evolutionary implications that can be drawn from this study. Ideally, a comprehensively-sampled global study using a combination of phylogenomic, population genomic, and morphological data should be undertaken to identify invasive lineages worldwide, clarify their relationships to native species, and enable a full taxonomic revision of *Lantana* sect. *Lantana*.

### Implications for biological control

We tested the concept of five flower-colour morphotypes employed by Australian biological control researchers to classify lantana. Our findings demonstrate that this concept fails to distinguish between different genetic clusters of potentially different provenance. Thus, identifying host background using flower colour alone might result in prospecting for biological control agents in the wrong locations, and/or deploying agents on the wrong hosts. The full varietal treatment of Smith & Smith (1982), which does identify distinct genetic clusters, has been under-utilised in biological control because it was found to be too difficult for practitioners to apply. Our study supports the development of improved identification tools for weed managers, e.g., field guides, and/or molecular diagnostics when field identification is challenging.

Variation in host preferences of some Australian lantana biocontrol agents aligns with our results. For example, host-specificity tests found that the leaf rust *Prospodium tuberculatum* could only establish on pink-flowered plants, but also that ∼25% of the pink-flowered plants tested were immune to it (Thomas et al., 2006). Susceptible hosts came from sites matching our predicted distribution of “Common Pink” (cluster A) while non-susceptible pink plants came from outside this range (Fig. 5), indicating that *P. tuberculatum* is specific to “Common Pink”. This supports the hypothesized Brazilian ancestry of this variety, since *P. tuberculatum* introduced to Australia was collected in Brazil (Thomas et al., 2006). Similarly, the flower-galling mite *Aceria lantanae* has generally failed to establish on pink-flowered lantana, except in northern Queensland (Murree 2018; 2023) within the predicted range of “Townsville Red-centred Pink” (cluster F). We therefore suggest “Townsville Red-centred Pink” is susceptible to *A. lantanae*, whereas “Common Pink” is not. The hypothesised provenance of the susceptible host (North America) again aligns with the agent’s introduction source (Mukwevho et al., 2017).

**Figure 5.**
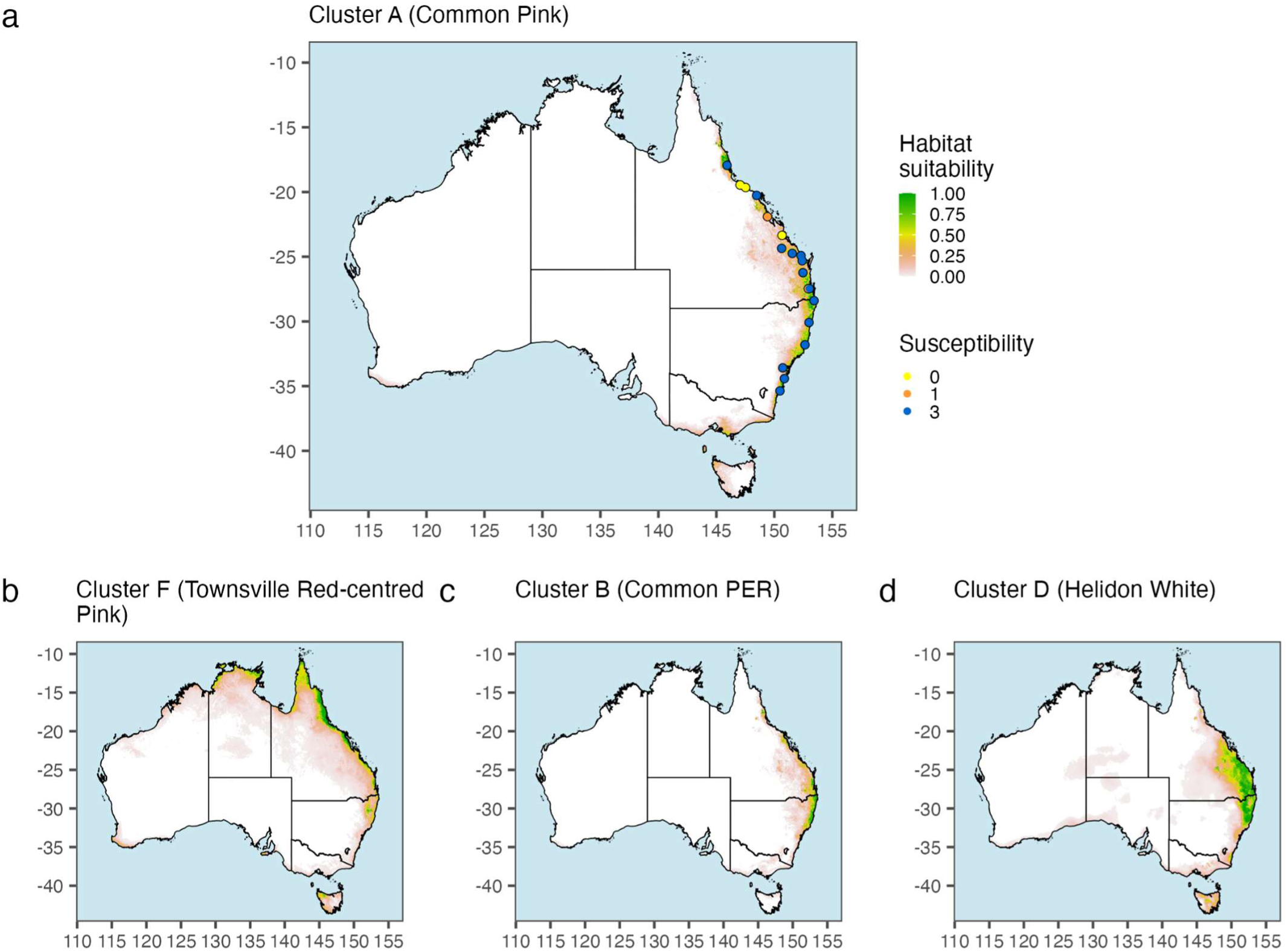
Predicted distributions of lantana clusters and host specificity testing. Predicted species distributions under current climatic conditions based on occurrence data for (A) cluster A (*Lantana camara s.l*. var. “Common Pink”), (B) cluster F (“Townsville Red-centred Pink”), (C) cluster B (“Common Pink-edged Red”), and (D) cluster D (“Helidon White”). In panel (A), collection coordinates of pink-flowered plants used for host specificity testing of the rust fungus *Prospodium tuberculatum*, introduced to Australia as a biological control agent, are overlaid. Points are colour-coded according to susceptibility (0 = not susceptible, 3 = highly susceptible; data from Thomas et al., 2006). Susceptible plants are located in areas of high predicted suitability for cluster A, while less susceptible plants are found in areas of lower predicted suitability.

Due to the presence of multiple distinct and divergent host genetic backgrounds within the invasive lantana complex in Australia, it is unlikely that any one biological control agent will provide effective control. Even so, biological control has an important role in integrated management approaches for lantana (Day et al. 2003a). To maximise effectiveness, different suites of agents will need to be tailored to different areas of lantana infestation where specific host varieties are dominant. Our findings provide a critical foundation for designing such customised biological control strategies. Additionally, by providing testable hypotheses of the native provenance of different genetic clusters of invasive lantana, we set the stage for further prospecting for new, more effective biological control agents.

### Implications for weed impact and risk assessment

Most ecological and economic impacts of invasive lantana in Australia can be attributed to the two most widespread varieties, “Common Pink” (cluster A) and “Common Pink-edged Red” (cluster B), which we predicted already occupy most of their suitable habitat. Orange- and red-flowered varieties (clusters C, E, G) were predicted to have limited suitable habitat nationally, but the accuracy of habitat suitability predictions for these rarer varieties may have been affected by lower numbers of available records (Elith et al., 2011). On the other hand, Maxent performs better with few records relative to other methods (Pearson et al., 2007) and our use of many environmental predictors should have improved accuracy in this case (Merow et al., 2013). In contrast, we predicted suitable habitat for “Helidon White” (cluster D) and “Townsville Red-centred Pink” (cluster F) far beyond their current distributions (Fig. 5b; Fig. S2). Comparing the distribution of “Helidon White” in our study (Fig. S2) with Smith & Smith (1982) reveals a ∼200km southward range expansion during the past ∼40 years, whereas the distribution of “Townsville Red-centred Pink” was unchanged. In our estimation, “Helidon White” is the lantana variety with greatest unrealised invasion potential in Australia.

Prior predictions of current and future suitable habitat for lantana in Australia have varied greatly depending on the modelling methods used (Adhikari et al., 2024; Goncalves et al., 2014; Taylor et al., 2012; Taylor & Kumar, 2013; van Oosterhout, 2004). These studies treated the complex as a single species. We showed that different varieties of lantana occupy different environmental niches, implying that better taxonomic resolution should improve the results of future modelling. Further, since soil moisture variables were the strongest predictors of distribution in all lantana varieties, these variables should be taken into account in future studies.

Invasive lantana in Australia is highly heterogeneous, and the different genetic clusters differ in their distributions, biological control agent susceptibility, and risk for future spread. They also differ in their potential impacts, e.g., toxicity to livestock (Smith & Smith, 1982). Thus, we argue that Australian weed policy should include scope to treat different lantana varieties differently, rather than blanket legislation for the entire species complex. Recognition of the different impacts by different varieties would enable more flexible and context-dependent risk assessment and management across jurisdictions, which would support more efficient mitigation of future impacts. In this context, a revised taxonomic treatment for invasive lantana is urgently needed.

## Conclusion

We showed that the evolutionary history and taxonomy of invasive Australian lantana is complicated, but we also clarified longstanding issues. We identified distinct genetic clusters corresponding to published varieties, likely descended from multiple progenitor species from different parts of the native range. The varieties “Common Pink” and “Common Pink-edged Red” are dominant in Australia, while “Helidon White” and “Townsville Red-Centred Pink” pose the greatest future invasion risk. Population genomic data over a large geographic area was the key to identifying genetic structure within invasive lantana, which had previously only been attempted using smaller-scale data and sampling. This project provides a framework for continuing to characterise lantana on a global scale, and a model for resolving invasive species complexes where management can be improved by better understanding of their diversity.

## Supporting information

sample metadata

## ACKNOWLEDGMENTS

We acknowledge the traditional owners of the lands throughout Australia on which this study was conducted. We thank the volunteers and colleagues representing multiple landcare, government, and research organisations who contributed samples to this study with permission (Terry Moody, Ailee Calderbank, Joseph Neilson, Jeff Harbrow, David Burgin, Russell Parr, Seamus Faithfull, Marion McCutcheon, Graeme Carrad, Maureen Magee, Renata Phelps, Jane Slack, Anne Boyden, Robyn Lamond, William DeGeer, Cheryl Moody, Robyn Hyde, Stuart Worboys, Kelli Murree, Barbara Waterhouse, Mark Gardener, Stephen McKenna, Miranda Sinott-Armstrong, Marcos Caraballo, Verônica Thode, Laura Frost, John Chau, Crystal Shin, David Giblin, Richard Olmstead, David Hembry, Katrina Dlugosch). Assistance with analyses was provided by Nunzio Knerr. We are grateful to Maurizio Rossetto and members of the Research Centre for Ecosystem Resilience, Botanic Gardens of Sydney for their advice and support. This study was funded by the Australian Government, Department of Agriculture, Fisheries and Forestry (activity ID: 4-FY9KQZ3).

## AUTHOR CONTRIBUTIONS

P.L-I., F.E-V., M.D., and J.J.L.R. designed the study. M.D., J.C., P.L-I., F.E-V., and J.J.L.R. conducted and coordinated the sampling. P.L-I., F.E-V., and E.M. analysed the data. P.L-I. and E.M. drafted the manuscript with major input from F.E-V, M.D., and J.J.L.R. All authors provided critical revisions and edits to finalise the manuscript.

## CONFLICT OF INTEREST STATEMENT

The authors declare no conflicts of interest.

### BIOSKETCH

The authors of this paper represent a range of perspectives and expertise, from multiple research programs and institutions based in Australia. The main interests of the authors include ecology, evolutionary biology, invasion biology, plant systematics, population genomics, and weed biological control. Several members of the research team are particularly interested in invasive plants, and have collectively authored dozens of publications on this topic over the past 30 years.

## DATA ACCESSIBILITY STATEMENT

The raw genotype matrices analysed in this study are available via Zenodo (https://doi.org/10.5281/zenodo.17224144) and the metadata are included as a supplementary file. Climate data (9 s resolution) are available from the CSIRO Data Access Portal at https://doi.org/10.25919/5dce30cad79a8 (Bio4-Bio32), https://doi.org/10.4225/08/5afa9f7d1a552 (EAA, EPI, WDI and WDX) and https://doi.org/10.4225/08/5b285fd14991f (CLY, SND, NTO, PTO, BDW and TWI3S).

NOTE genotype data are currently restricted and will be made accessible upon publication of manuscript.

