## supporting tables and figures for "Tangled evolutionary history: genetically divergent taxa and hybrids characterise lantana invasions in Australia"

### SUPPLEMENTARY FIGURES & TABLES

###### Table S1. Details and metadata for samples used in this study.

[.csv file attached separately]

| ***Genetic cluster*** | ***AUC*** | ***Kappa*** | ***MTSS*** | ***Environmental predictors*** |
| --- | --- | --- | --- | --- |
| Cluster A | 0.99 | 0.65 | 0.2 | *B28, EAA* |
| Cluster B | 0.99 | 0.95 | 0.06 | *B28, EAA* |
| Cluster C | 1.0 | 0.95 | 0.95 | *B28, PTO* |
| Cluster D | 0.98 | 0.85 | 0.52 | *WDI, B31* |
| Cluster E | 1.0 | 0.93 | 0.93 | *EAA, B31* |
| Cluster F | 0.99 | 0.92 | 0.32 | *B32, EAA* |
| Cluster G | 1.0 | 0.99 | 0.98 | *B32, B06* |

###### Table S2. Species distribution model performance statistics for all genetic clusters of Australian lantana.

Area under the Received Operating Characteristic curve (AUC), maximum of Cohen’s Kappa (K) threshold, and Maximum training sensitivity and specificity logistic threshold (MTSS). We also show the two most important environmental predictors for each genetic cluster’s SDM found by MaxEnt in the column called ‘Environmental predictors’.B06: Min temperature of coldest week (°C); B28: Annual mean moisture index; B31: Moisture index seasonality (C of V); B32: Mean moisture index of wettest quarter; EAA: Annual total actual evapotranspiration; PTO: Total Phosphorus; WDI: Minimum monthly atmospheric water deficit (precipitation - potential evaporation).
Table S3. Summary statistics of individual heterozygosity and inbreeding for each genetic cluster.

Median observed heterozygosity (*H_O_*) and inbreeding coefficient (*F_IS_*) of individuals within each cluster, with standard deviation (SD) shown. Sample size (*n*) indicates the number of individuals per cluster. Data presented here is the same as in Fig. S3.

| Cluster | *n* | *H_O_* (median ± SD) | *F_IS_* (median ± SD) |
| --- | --- | --- | --- |
| A | 115 | 0.015 ± 0.004 | 0.923 ± 0.022 |
| B | 82 | 0.003 ± 0.003 | 0.986 ± 0.017 |
| C | 12 | 0.153 ± 0.02 | 0.198 ± 0.103 |
| D | 46 | 0.12 ± 0.022 | 0.371 ± 0.114 |
| E | 13 | 0.151 ± 0.034 | 0.208 ± 0.177 |
| F | 39 | 0.019 ± 0.008 | 0.9 ± 0.042 |
| G | 11 | 0.06 ± 0.045 | 0.688 ± 0.234 |
| Unclustered Australian | 206 | 0.108 ± 0.056 | 0.434 ± 0.294 |
| Native range | 128 | 0.026 ± 0.031 | 0.863 ± 0.165 |

###### Table S4. Adjusted p-values from post hoc pairwise Wilcoxon tests (Bonferroni correction) comparing individual heterozygosity (H_O_) between genetic clusters.

The Kruskal-Wallis test showed a significant difference in *H_O_* among clusters (χ² = 403.38, df = 8, *p* < 2.2e-16). Values represent adjusted p-values for pairwise comparisons, with significant differences (*p* < 0.05) highlighted.

|  | A | B | C | D | E | F | G | Unclustered Australian |
| --- | --- | --- | --- | --- | --- | --- | --- | --- |
| B | 0.000 |  |  |  |  |  |  |  |
| C | 0.000 | 0.000 |  |  |  |  |  |  |
| D | 0.000 | 0.000 | 0.001 |  |  |  |  |  |
| E | 0.000 | 0.000 | 1.000 | 0.022 |  |  |  |  |
| F | 0.260 | 0.000 | 0.000 | 0.000 | 0.000 |  |  |  |
| G | 0.000 | 0.000 | 0.020 | 1.000 | 0.055 | 0.000 |  |  |
| Unclustered Australian | 0.000 | 0.000 | 0.016 | 1.000 | 0.071 | 0.000 | 1.000 |  |
| Native range | 0.000 | 0.000 | 0.000 | 0.000 | 0.000 | 0.029 | 0.011 | 0.000 |

###### Fig. S1. Genome size estimates

Estimates of genome size (DNA content) in picograms from flow cytometric analysis of nuclei extracted from 14 wild-sampled individuals of two lantana morphotypes.
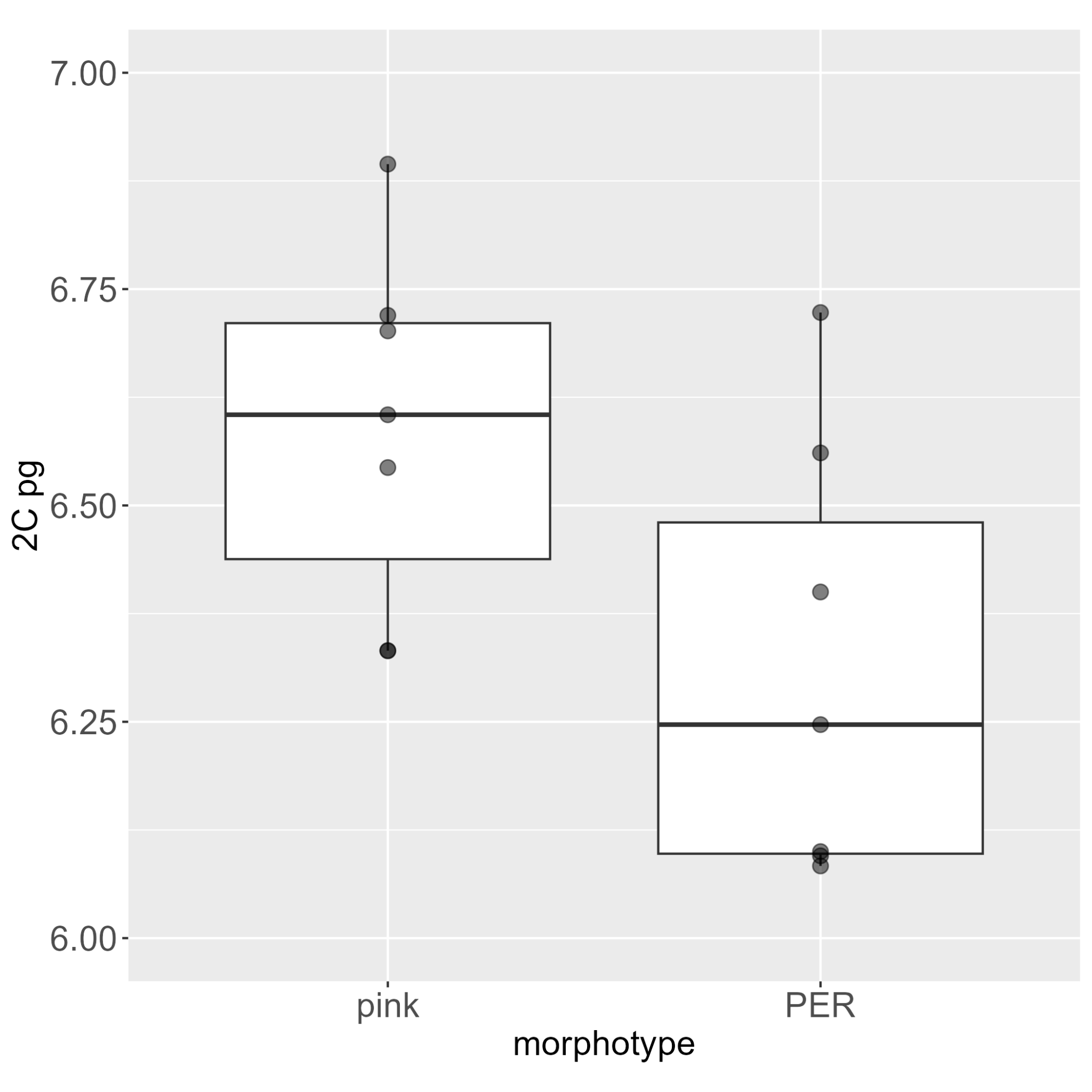

###### Fig. S2. Genetic cluster maps

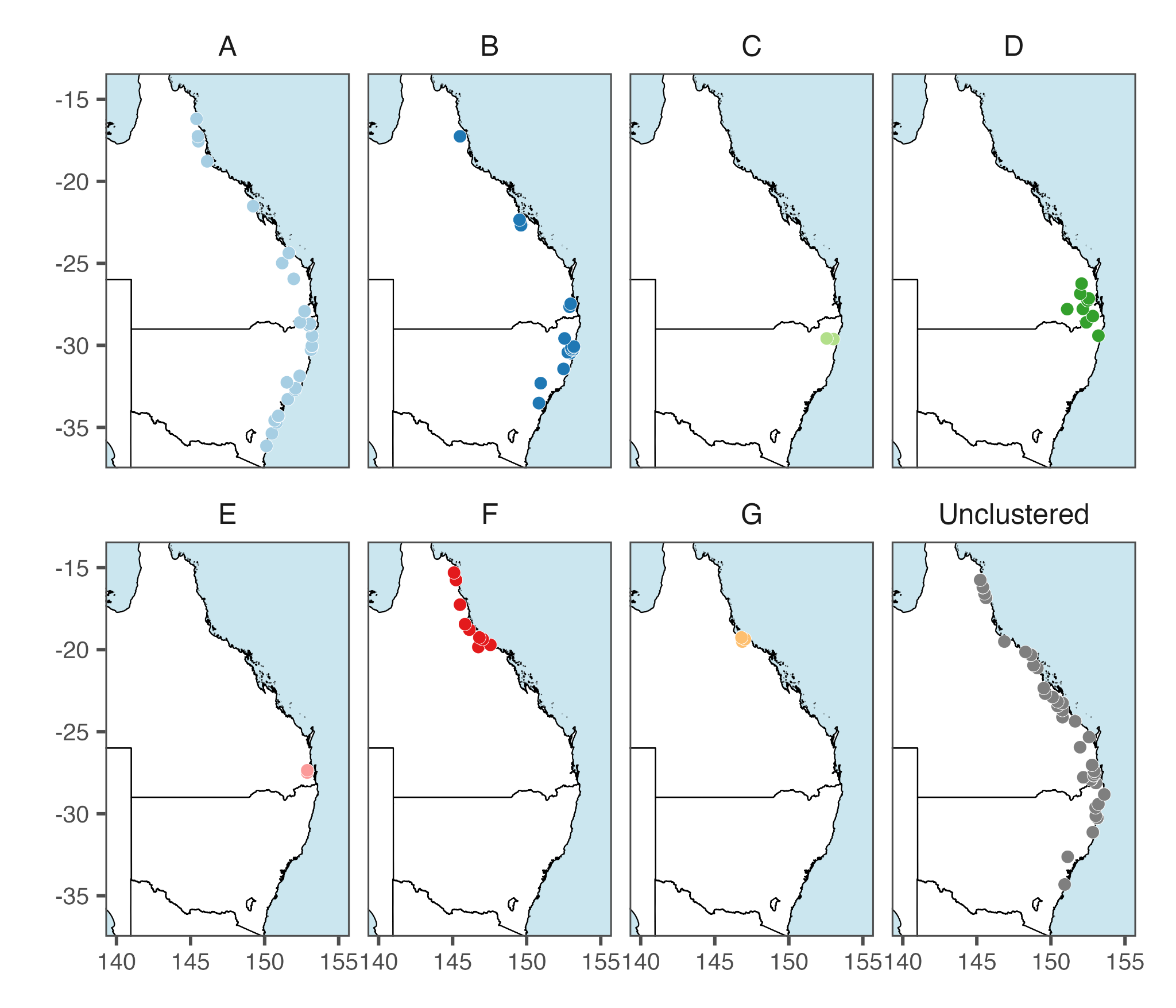

Geographic distribution of samples assigned to genetic clusters based on population analysis. These occurrence points were used to construct the HSMs.

###### Fig. S3. Individual Heterozygosity

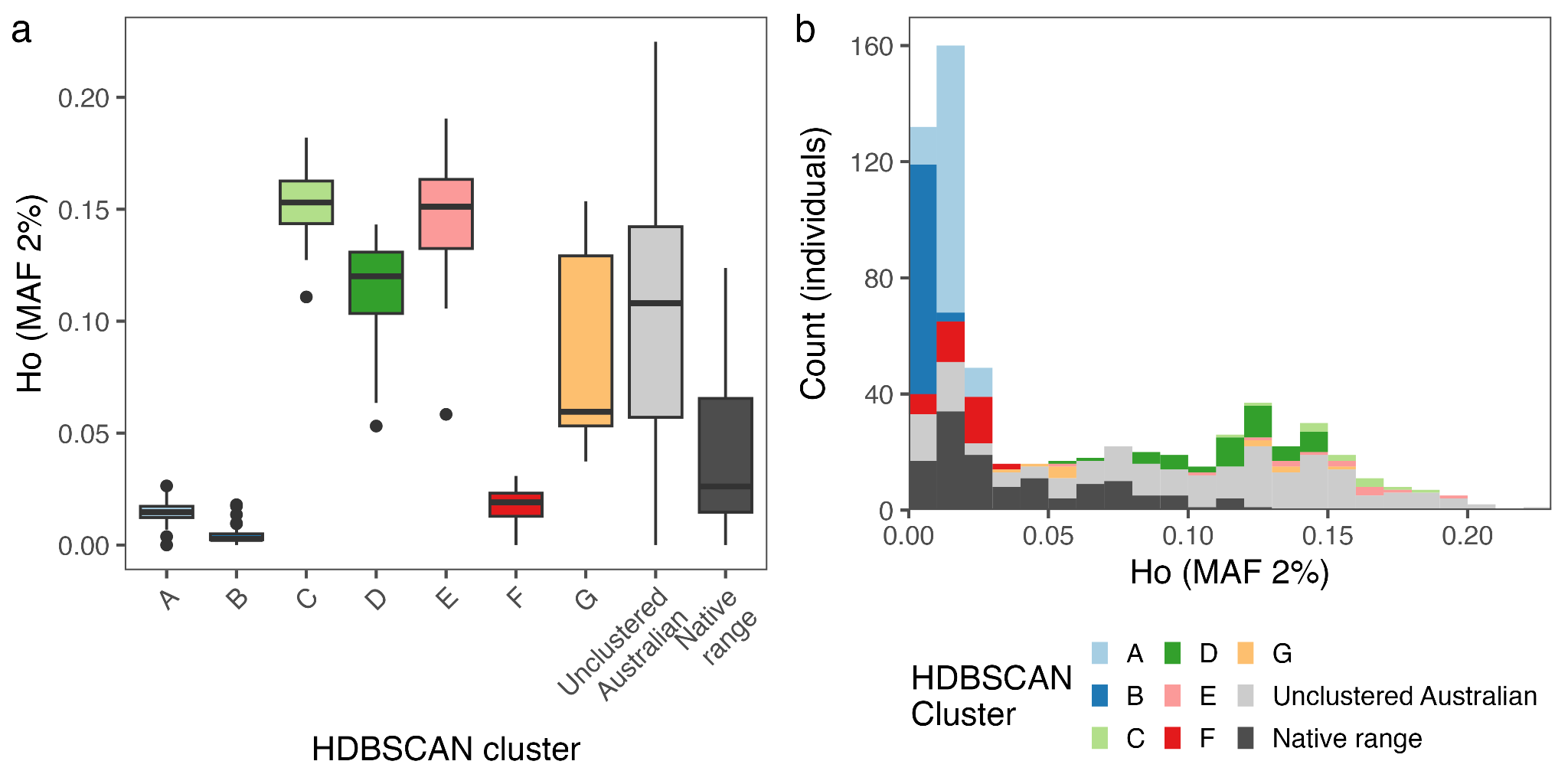

Heterozygosity at loci with a minimum minor allele frequency (MAF) > 2%. (A) Boxplot displaying heterozygosity by genetic cluster. (B) Histogram showing the distribution of the proportion of heterozygous loci across all samples.

###### Fig. S4. Extended LEA snmf results

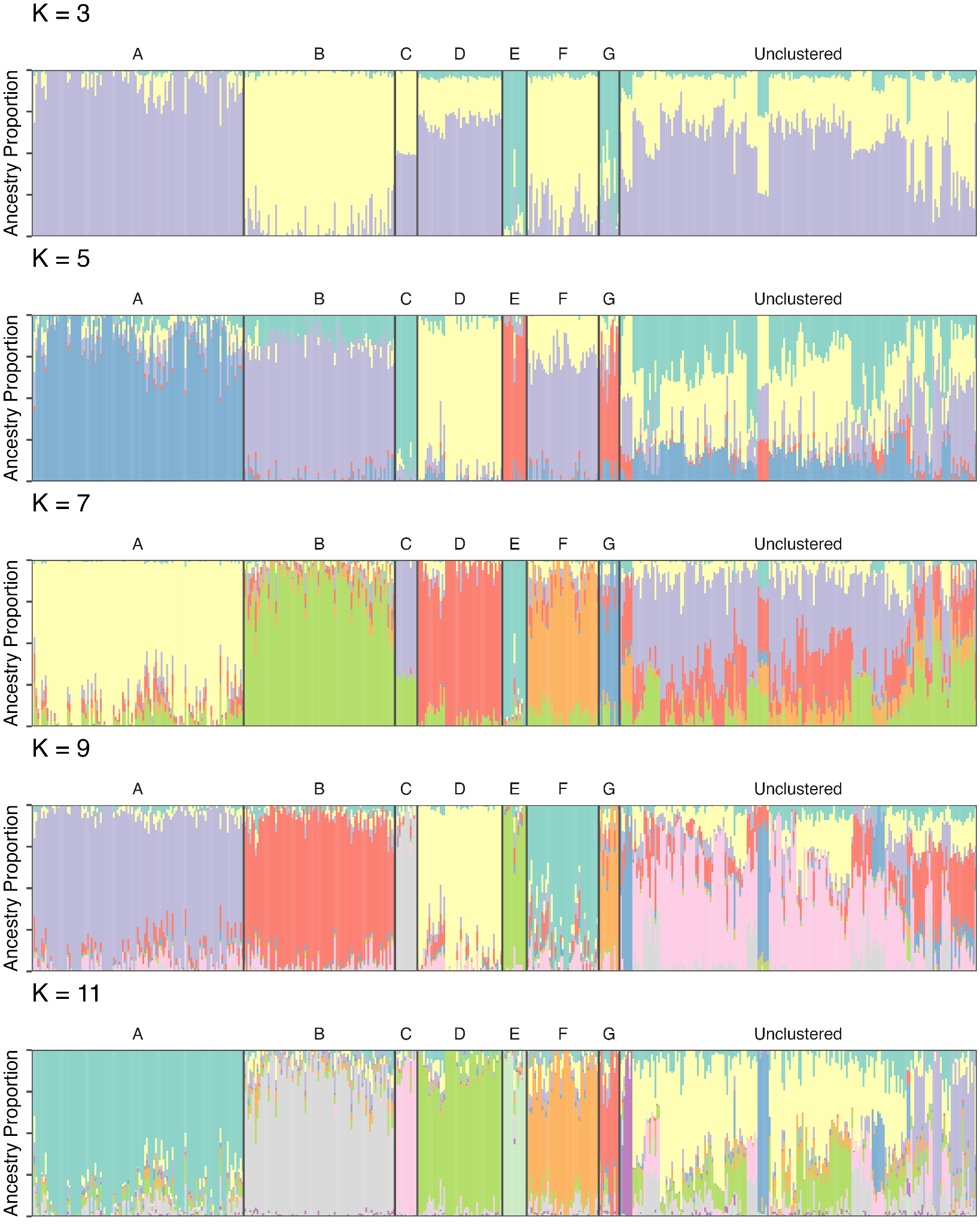

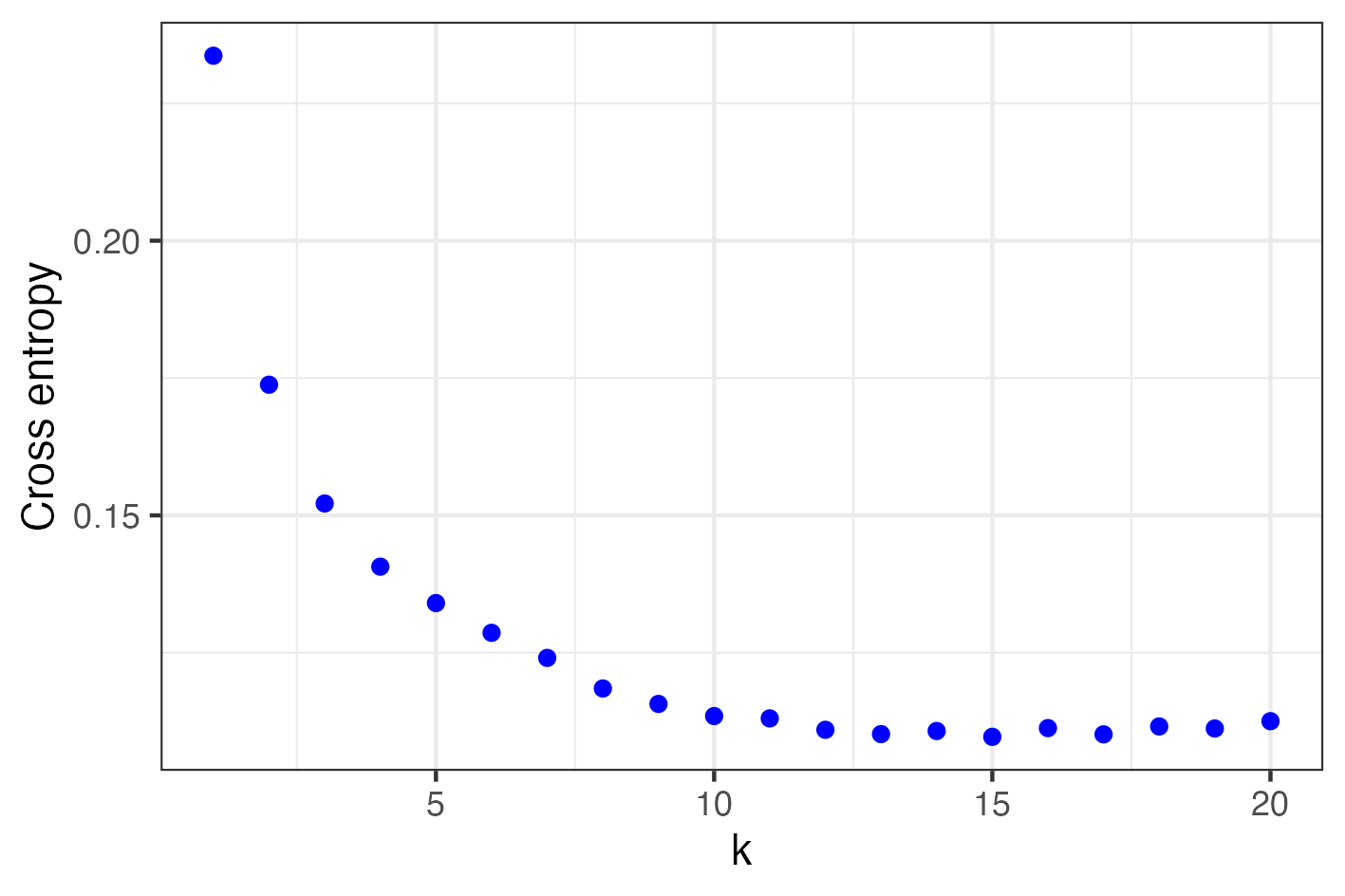
Ancestry estimation by LEA snmf for K = 3, 5, 7, 9, and 11 on eastern Australian lantana, grouped by genetic clusters (top); cross-entropy plot for K = 1-20 (bottom).

###### Fig. S5. Putative hybrid results

Hybridization evidence between “Common Pink” and “Common Pink-Edged Red” *Lantana camara*, based on 33 individuals from four sites (two with uniform flower colour and two with mixed/intermediate colours).

(a) Plot of the first principal component axis (PC1) vs. observed heterozygosity (4,986 loci), with putative hybrids clustering in the center. PC1 is the same as in Fig. 2a.

(b) Heatmap of loci fixed in “Common Pink” and “Common Pink-Edged Red” genetic clusters, with yellow = fixed reference alleles, blue = fixed alternate alleles, green = heterozygous loci, and white = missing data. Putative hybrids are predominantly heterozygous (green) at sites which are fixed for non-hybrid individuals.

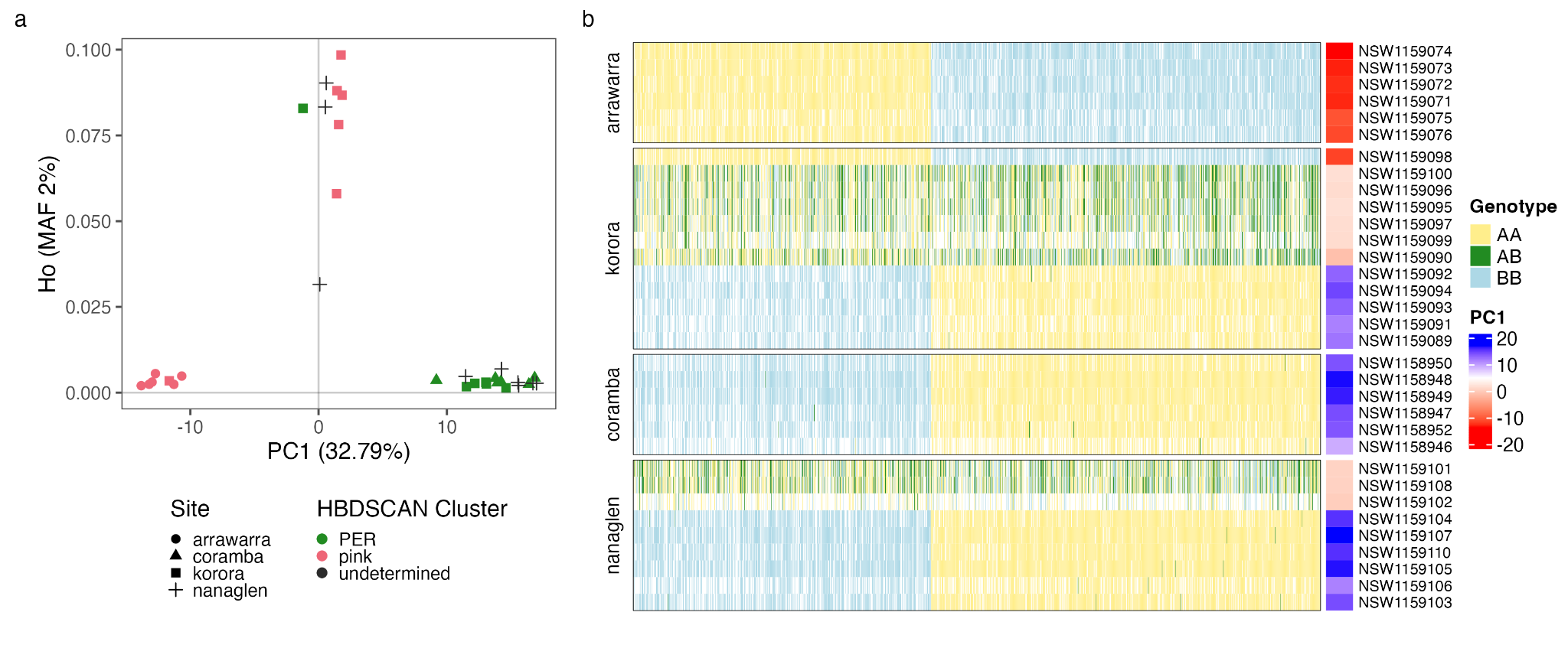
For the allele heatmap, data was filtered to retain relevant samples, remove loci with <98% reproducibility, exclude fixed loci, and keep one SNP per DArT tag. Alternatively fixed SNPs between parent groups were identified with a 5% allele frequency leniency (fixed >95%), and loci with >80% missing data were excluded, resulting in 1,455 loci.

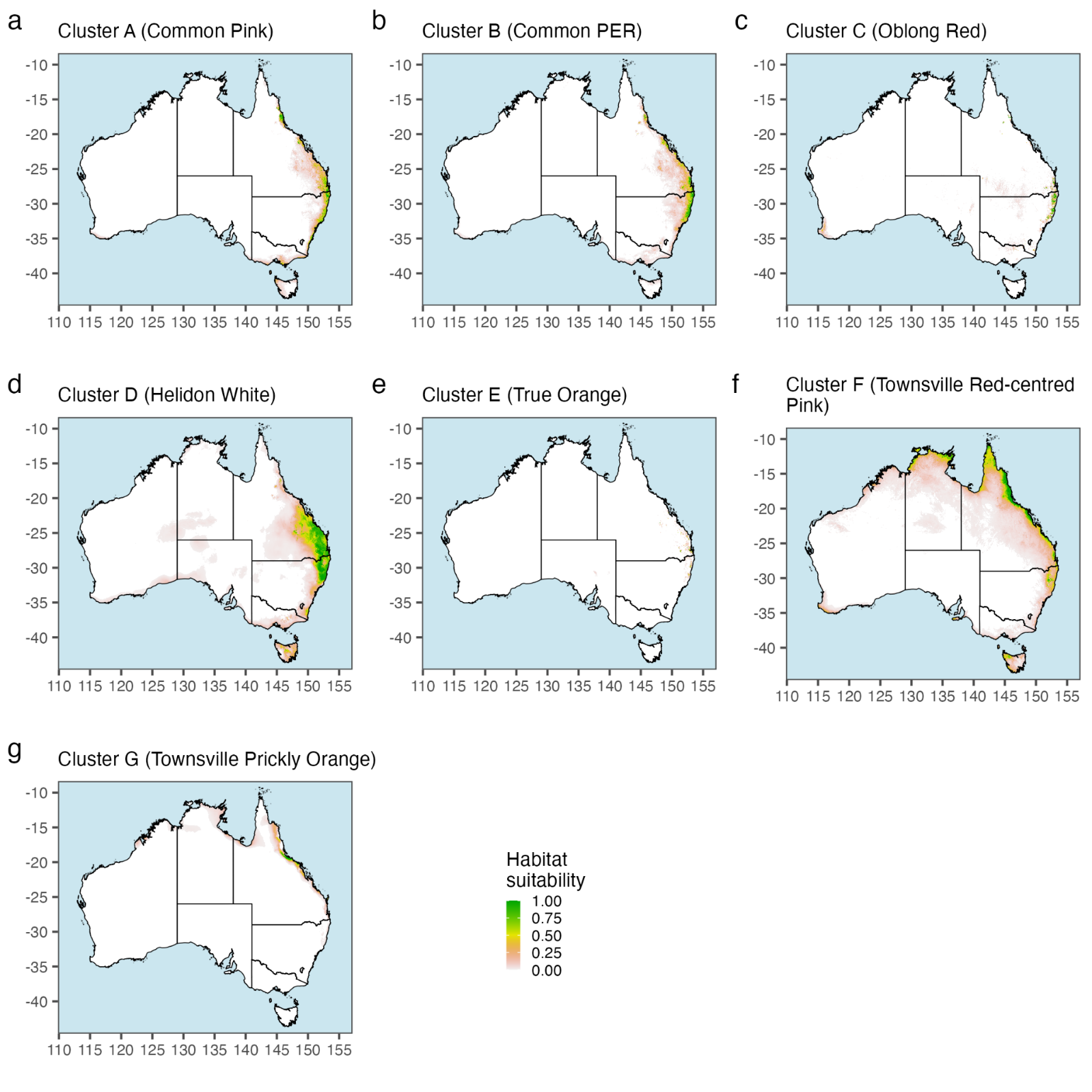

###### Fig. S6. Species distribution models for all genetic clusters.
